# A 1D Model Characterizing the Role of Spatiotemporal Contraction Distributions on Lymph Transport

**DOI:** 10.1101/2022.09.26.508881

**Authors:** Farbod Sedaghati, J. Brandon Dixon, Rudolph L. Gleason

**Affiliations:** The George W. Woodruff School of Mechanical Engineering, Georgia Institute of Technology, Atlanta, GA; The Wallace H. Coulter Department of Biomedical Engineering, Georgia Institute of Technology, Atlanta, GA

**Author notes:** <u>Corresponding Author:</u> Rudolph L. Gleason, Jr., Ph.D. The George W. Woodruff School of Mechanical Engineering, The Wallace H. Coulter Georgia Tech/Emory Department of Biomedical Engineering & The Petit Institute for Bioengineering and Bioscience, Georgia Institute of Technology, 387 Technology Circle 05, Room 205, Atlanta, GA 30313.

**Keywords:** lymphatic system, lymph flow, computational model, spatiotemporal contraction, wall shear stress

## Abstract

**Background:** Lymphedema is a condition in which the two primary functions of the lymphatic system, immunity and lymph transport, are compromised. Computational models of lymphatic function to characterize lymph transport have proved useful in quantifying changes in lymph transport in health and disease; however, much is unknown regarding the regulation of contractions throughout a lymphatic network. The purpose of this paper is to study the role of pacemaking cells and contractile wave propagation on lymph transport using a 1-D mathematical model.

**Method:** A 1D fluid-solid modeling framework with constitutive relationships were employed to characterize the interaction between the contracting vessel wall and the lymph flow during contractions. The time-space distribution of contraction waves along the length of a three-lymphangion chain, was determined by prescribing the location of pacemaking cells and parameters that govern the contractile wave propagation, with the total contractile response at each location determined as the summation of the local electrical signals.

**Results:** Spatiotemporal changes in radius and lymphangion ejection fraction from our illustrative simulations mimic well values reported in isolated lymphatics from the wild-type (WT) and Smmhc-CreER^T2^;Cx45^fx/fx^ knock-out (KO) reported in the literature. The flow rate is 5.43 and 2.45 *μL/hr* in the WT and the KO (average) models, respectively. The average and the peak WSS in the WT model are 0.08 and 4 *dyn/cm*^2^ and −0.03 and 0.87 *dyn/cm*^2^ in the KO (average) models, respectively.

**Conclusions:** The factors that govern the timing of lymphatic contractions remain largely unknown; but these factors likely play a central role in health and disease. This modeling framework captures well lymphatic contractile wave propagations and how it relates to lymph transport and may motivate novel assays and experimental endpoints to evaluate subtle changes in lymphatic pumping function in the development and progression of lymphedema.

## INTRODUCTION

The lymphatic system transports the interstitial fluid back to the venous system (usually) against an adverse pressure gradient. Lymph is collected through primary (initial) lymphatics, gets transported through contractile and collecting (secondary) lymphangions primarily via orchestrated lymphatic muscle cell contraction, then through lymph nodes and ducts into the circulatory system. The primary lymphatic vessels absorb the interstitial fluid by communicating with the surrounding interstitium via primary lymphatic valves and transport the generated lymph. To achieve this goal, secondary lymphatics contract autonomously to drive the lymph through one-way valves. This autonomous contractile behavior is governed by numerous factors including mechanical stretch of lymphangions, the chemical microenvironment, electrical conductivity, amongst many other factors. Dysfunction of the lymphatic system leads to lymphedema, a condition characterized by swelling and accumulation of lymph in a particular limb or region. Disruption of lymph transport in lymphedema caused by deviations in contractile factors is to be addressed in this study.

Computational models have been used extensively to study any biological systems including plants, animals, and human physiology, specifically in developing any abnormality, disease, or injury^1–6^. Moreover, they have proven useful in characterizing lymphatic contractile behavior and lymph transport. However, much is unknown with regard to the regulation of contractions, particularly the role of propagation of action potentials in regulating pump action. There are some mathematical approaches to study the lymphatic mechanics, but so far, lumped-parameter models have been extensively used to simulate lymph transport. Also, to best of our knowledge, no one has implemented contractile parameters in the computational models. In such models, the pressure gradient and the contraction frequency of a lymphatic vessel are input to the solver and the transmural pressure and flow are calculated. The drawbacks of such models are that the pacemakers cannot specifically be located, the contraction propagation speed is not applied, and the pressure/flow waves along a lymphangion are missing^7^. Another drawback of the lumped-parameter model is that the nonlinearity of the balance laws is not addressed in the algebraic equations. For example, Bertram et al. showed that the pressure evenly ramps up along a lymphatic chain which may not be the case^8^. Contraction coordination of multi-lymphangions is an interesting aspect that is not accessible in lumped-parameter models focusing on single lymphangions^9^. Also, the lymphatic function is highly disturbed in damaged lymphatic vessels where both the contractile and the mechanical properties are compromised^10^. To address these issues, a 1-D discretized model would help quantify the physical fields that are time and space dependent. In a recent 1-D study, Elich investigated how lymph flow changes in different contraction scenarios but they have not implemented contraction metrics^11^.

Lymphedema is usually accompanied by growth and remodeling of lymphatic vessels which most likely happens through flow-contraction coupling mechanisms. Such coupling mechanisms have proved in previous experimental studies where the contraction outcomes (amplitude and frequency) are monitored in a controlled mechanical environment (stretch, luminal pressure, and wall shear stress)^12–15^. Also, it is proved that the imposed flow inhibits the active pumping in lymphatics^16^.In a study, they included contraction-WSS coupling which again due to the lumped formulation, the WSS distribution along a vessel where pacemakers are located is not included^17^. These mechanisms are not accurately applicable in lumped-parameter models. In the microscopic view, chemical concentrations (NO, Ca) at the cellular levels are the hidden chain between the mechanical environment and the contractility. A few studies have implemented the chemical and the pressure/wall shear stress couplings in the mathematical models, but due to uncertainty and the lack of experimental measures, they are not applied in the current study^18,19^. Another aspect is including the initial lymphatics and study Guyton Suction Theory in absorbing lymph from the interstitium which is a challenging question^20^. Compromised lymph absorption in lymphedema is an interesting topic that has been addressed in a few studies and will be added to the model later^21–23^.

As the main controller of lymphatic contractility, the activation function has been extensively explored in the mathematical simulations. Researchers usually use trigonometric functions with some time delay between lymphangions to prescribe contractions along a lymphatic chain^24,25^. Not using experimental pacemaking metrics and getting conflicting results are the main challenge in prescribing the activation function in lymphatics^26^. For example, in one of the first lymphatic experiments, they realized that a lymphangion may start contracting while adjacent ones are almost in the diastole phase and so contractions are not coupled^27^. Contrary, it was shown that adjacent lymphangions are electrically coupled meaning that any disturbances in a pacemaker affects nearby contractile activities^28^. Additionally, there is an ongoing debate that the maximum pumping occurs when there is no time delay between lymphangions^29^, while others assume that if lymphangions contract respectively, they generate a stronger propulsion^25^. Or McHale et al. suggested that antegrade pumping is more efficient^30^, while Venugopal et al. believed that the coordination of lymphangions and the direction of the contraction propagation minimally affect the pumped flow^29^. Based on the experimental observations, the delay and the opening duration of secondary valves are highly dependent on the activation function^31^. Or the mean lymph flow rate almost doubles by slightly changing the relaxation time in diastole^32^. Therefore, there is a pressing need to explore the mechanics of contractions and lymph transport more in details.

Therefore, the purpose of this paper is to develop a computational model to study the mechanics of the lymphatic system and quantify how abnormal wave propagation compromises the pumping efficacy. So, this study aims at constructing a 1-D mathematical model that is based on fluid-solid governing equations and utilizes a general activation function based on experimental contractility data.

## METHODS

Motivated by available experimental data in the literature^33^, we will model a lymphatic vessel that consists of three lymphangions and four secondary valves (***Figure 1***). The inlet and outlet pressures are prescribed as 3.5 and 4 *cmH*_2_*O*, respectively, for all illustrative simulations.

**Figure 1.**
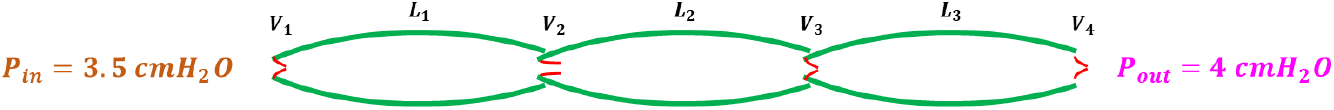
A schematic representation of the modeled lympahtic chain

### Theoretical Framework

#### 1D Fluid Mechanics Framework

Each lymphangion is modeled as a tapered, elastic tubes in a one-dimensional pulsatile fluid flow framework. Conservation of mass and the balance of linear momentum require that (Equation (1)),

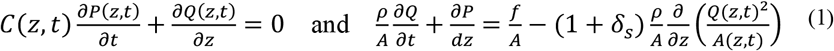

where *A*(*z, t*) is the instantaneous arterial lumen cross-sectional area, *P*(*z, t*) is the luminal pressure, *Q*(*z,t*) is the volumetric flow rate, and *C*(*z, t*) = *∂A*(*z,t*)/*∂P*(*z,t*) is the area compliance; *z* is the direction along the vessel axis, *t* is time, *ρ* is the blood density, *f* is the frictional force per unit length. We prescribe the velocity profile (Equation (2)), *u_z_*(*r, z, t*), at any instant *t* and location *z* as

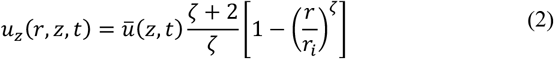

Where *ū*(*z, t*) = *Q*(*z, t*)/*A*(*z, t*) is the mean velocity, *ζ* is a constant that governs the shape of the velocity profile, r is the radial coordinate, and *r_i_* is the lumen radius. Using the Naiver-Stokes equations and integrating Equation (2) the viscous force is calculated as (Equation (3)),

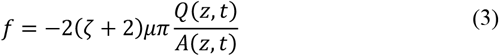

Where *μ* is the apparent lymph viscosity. This framework is explained in details in our previous study^4^.

#### Solid mechanics

To proceed, Equations (1) require information on the cross-sectional area, *A*(*z, t*), and compliance *C*(*z, t*) at each time (*t*) and location (*z*), given applied loads (*P*(*z, t*)) and activation state along the lymphangion wall. To characterize the wall behavior, we use data from Davis et al. (***Figure 2***), which represent the ‘pre-twitch’ and ‘peak-twitch’ mechanical states^31^. Pre-twitch represents the dilated configuration when muscle cells are fully relaxed and the peak-twitch state represents the configuration when muscle cells are at their peak contractile state over the pumping cycle.

**Figure 2.**
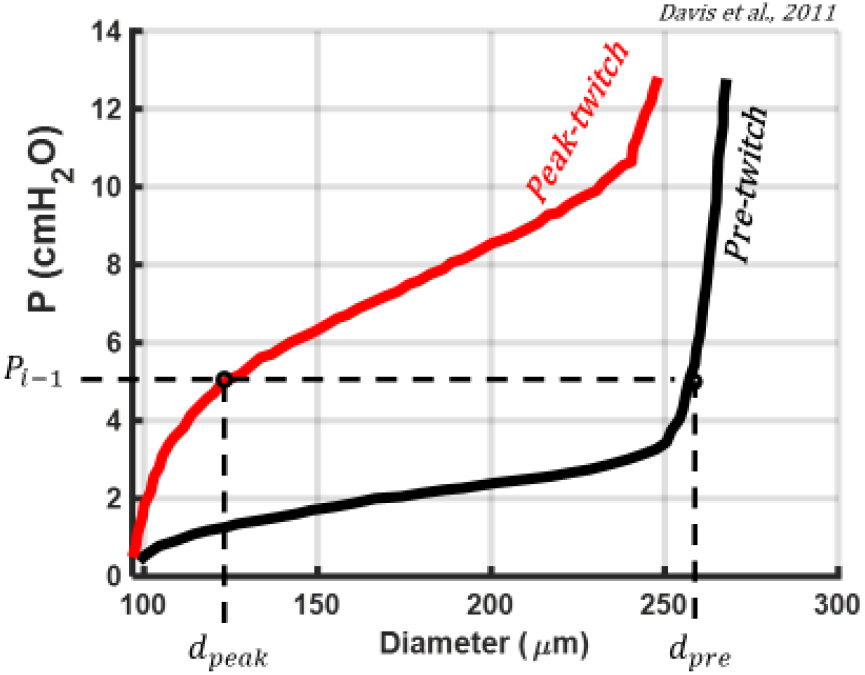
Pressure-diameter data during a contraction cycle of rat mesentery

To determine the diameter for a given pressure and level of muscle activation we employ a weighted-average of pre- and peak-twitch diameters where the weight function is the general activation function (Equation (4)); namely,

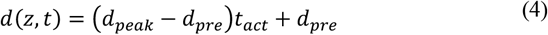

*d_pre_* and *d_peak_* are pre-stretch and peak-stretch diameters at pressure from the previous time step, and *t_act_* is the contraction parameter varying between 0 and 1. In contrast to using known constitutive laws for the active and passive mechanical response and solving the highly non-linear inverse solution of geometry (i.e., diameter), given applied loads (i.e., pressure) and active and passive material properties, Equation (4) allows one to approximate the diameter at each time point and location, with less computational cost and similar accuracy.

Area compliance may be expressed as (Equation (5)),

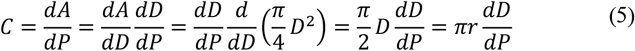

Thus, at pre- and peak-twitch contraction states (Equations (6)(7)),

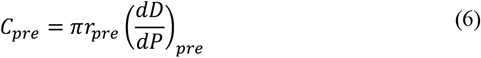

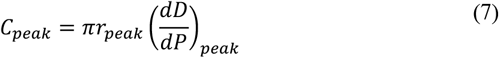

*C_pre_* and *C_peak_* are the pre-stretch and peak-stretch compliances at *r_pre_* and *r_peak_*, respectively. Also, (*dD/dP*)_*pre*_ and (*dD/dP*)_*peak*_ are the slope of the pressure-diameter curves. To find the pre-stretch and peak-stretch compliances we need to find the corresponding radius based on the pressure from the previous step as well as the derivative of the pressure-diameter curves. To approximate the compliance, we let (Equation (8))

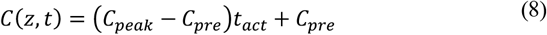

#### Valve mechanics

Lymphangions are defined as a piece of lymphatic vessel that is bounded between two pairs of leaflets that work like one-way valves and help with forward propulsion of lymph in the network. Due to the small sizes of the vessel, it is not easy to extract a valve and measure the mechanical properties of the leaflets. So, there are some experimental studies that have quantified the mechanics of secondary valves under different pressure gradient. Almost all these studies, propose a sigmoidal-type of curve for pressure vs. lymph flow. In this study, we use a valve behavior introduced by Bertram et al. (Equation (9))^25^.

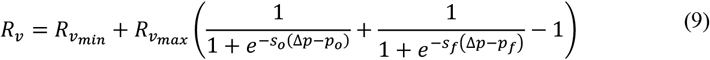

The parameters introduced in Equation (9) are provided in ***Figure 3***.

**Figure 3.**
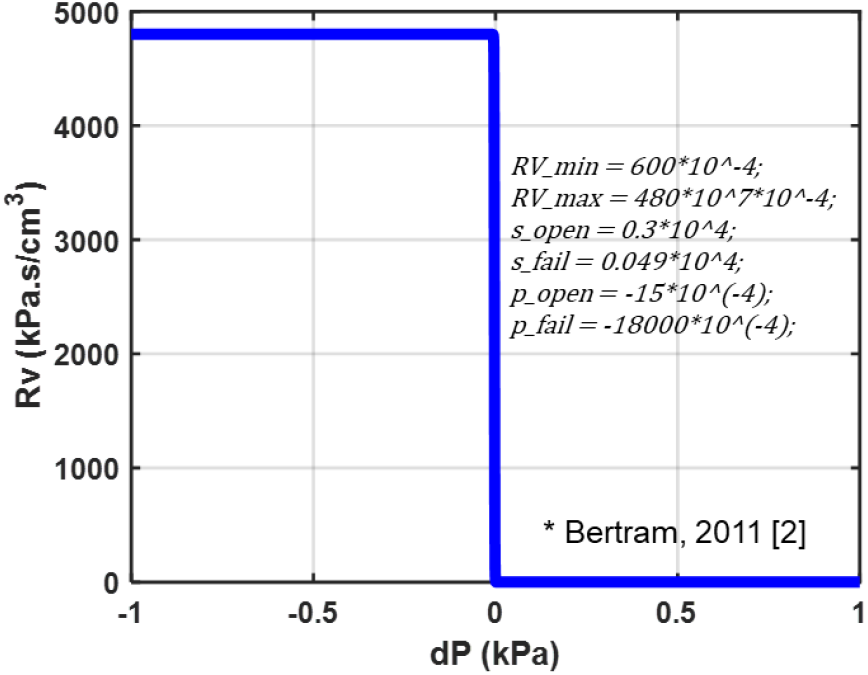
Pressure-valve resistance sigmoidal behavior

### Spatiotemporal activation function

One of the critical features in lymphatic contractility is pacemaking activity. Many studies have showed that there are some specific spots in a lymphatic vessel at which the contractions are generated. These locations are called pacemakers or pacemaking sites. Besides locating these cells, some metrics are usually measured to study the contractions such as contraction propagation speed, pacemaking distance, etc^34,35^. From these studies, we realize that contractions demonstrate a wave behavior which explains why a one-dimensional model is required to study the lymphatic system. Based on an experimental study by Zawieja et al., contractions in a lymphatic network propagates both temporally and spatially with a certain propagation speed^27^. Also, these waves have shown to transmit between adjacent lymphangions via gap junctions. This idea will be used as the core concept of the model in which the driver of the mechanical system is an activation parameter. As the key element of this model, an activation function dictates how each node in the network contracts. To prescribe contractions, researchers have used trigonometric functions that has a few disadvantages like they are not similar to the real activation signal, pacemaking parameters cannot be applied, and they are not applicable in 1-D models. To overcome these challenges, a 2-D Gaussian function is used in which the pacemaking parameters are implemented. In this function, the center of a distribution is placed at a pacemaker site (*μ_x_*) when it is supposed to generate a signal (*μ_t_*). The temporal standard deviation (σ_t_) is correlated to how rapidly the contraction wave decays over time. The physical interpretation of this parameter could be how fast ion channels, gap junctions, etc. get excited to transport the ions and generate the force. The spatial standard deviation (*σ_x_*) is correlated to how far a contraction wave travels in space. In other words, this parameter mimics cellular resistance and how easy the cells pass the contraction wave.

Below is a general form of a bivariate Gaussian equation (Equation (10)):

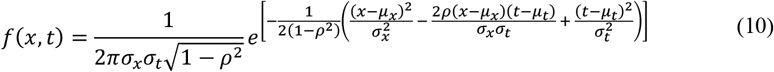

where *μ_x_* is location of the pacemaker that is determined by pacemaking distance and *μ_t_* represents the temporal location of the pacemaking signal. *σ_x_*, *σ_t_*, and *ρ* are the spatial, temporal, and correlated standard deviations. To prescribe them, we assume that contractions must decay between any two pacemakers in a contraction period timeframe. Spatially, there must be two decays between any two pacemakers. Mathematically one decay is almost 4 standard deviations meaning 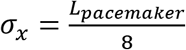. Similarly, there must be one decay in a contraction period resulting in the standard deviation of 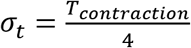. *ρ* which is called “correlation”, represents the contraction speed and shows how the distribution is skewed in the time and the space domains. The minimum of *ρ* is 0 meaning the contraction propagation speed is high and the distribution is not skewed. From the definition of Jacobian of the covariance matrix, the maximum limit of *ρ* is 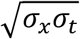. This happens when the propagation speed is low and the distribution is highly skewed in the space-time domains. ***Figure 4*** is a schematic representation of the standard deviations.

**Figure 4.**
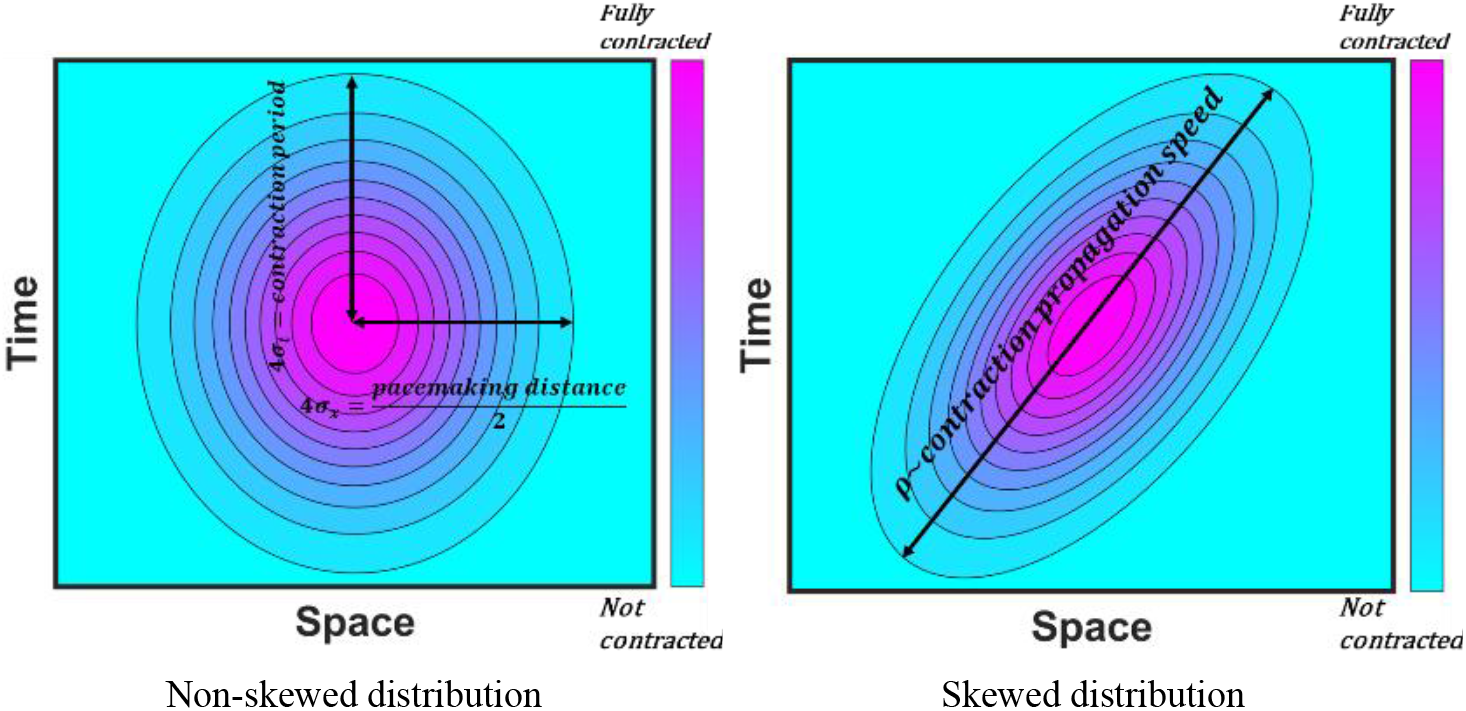
Representation of three standard deviatioans of Gaussian function and relationships with pacemaking metrics

In order to construct the overall activation distribution of a lymphatic chain, we use the concept “Summation” that has widely been investigated in neurophysiology. Based on this concept, the resulting signal is created from multiple simultaneous (spatial) and repeated (temporal) signals. This is applied in our model to add up Gaussian functions generated at pacemakers to construct the overall spatiotemporal contraction distribution of a lymphatic network (***Figure 5***). More specifically, the activation parameter of each node is the normalized algebraic summation of all signals at that node (Equation (11)).

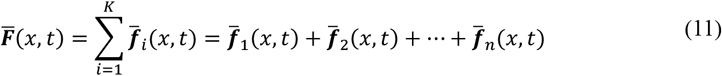

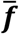 is a 2-D Gaussian function located at each pacemaker, *K* is the number of pacemakers, and 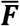 is the general activation function that covers the whole time-space domains. Equation (12) calculates *t_act_* for each node at each time step.

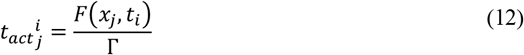

**Figure 5.**
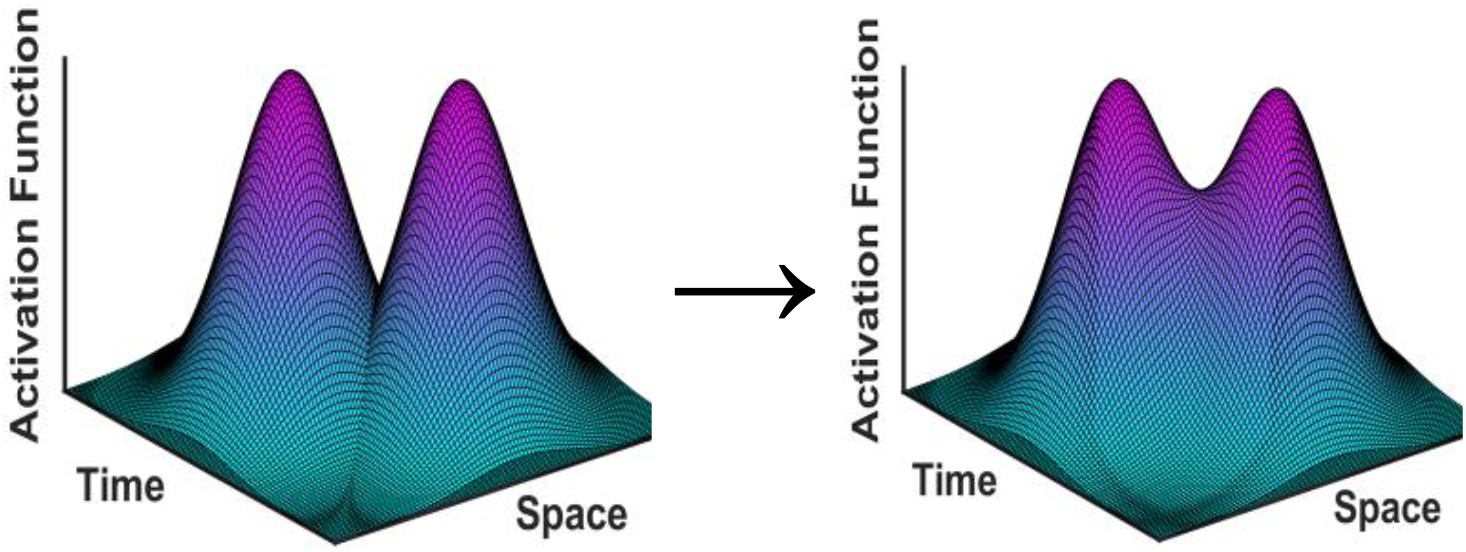
A representation of how “Summation” is used in the model

Γ is the maximum activation parameter of all nodes during a contraction period.

### Mathematical solver

An implicit finite difference scheme was devised to solve the governing equations with a given vascular geometry and material properties. The collecting lymphangions are discretized with dz of 50 *μm* and the time domain into time steps of 5 *msec*. At each time step the general matrix of unknown pressures and flows are calculated. As a metric to verify robustness of the solver, conservation of mass is verified during each contraction cycle, meaning that the cycle-averaged flow entering the first lymphangion is unchanged along the chain. ***Table 1*** shows a list of assumptions of the model. The discretization method is explained in details in ***Appendix. A***.

**Table 1.**
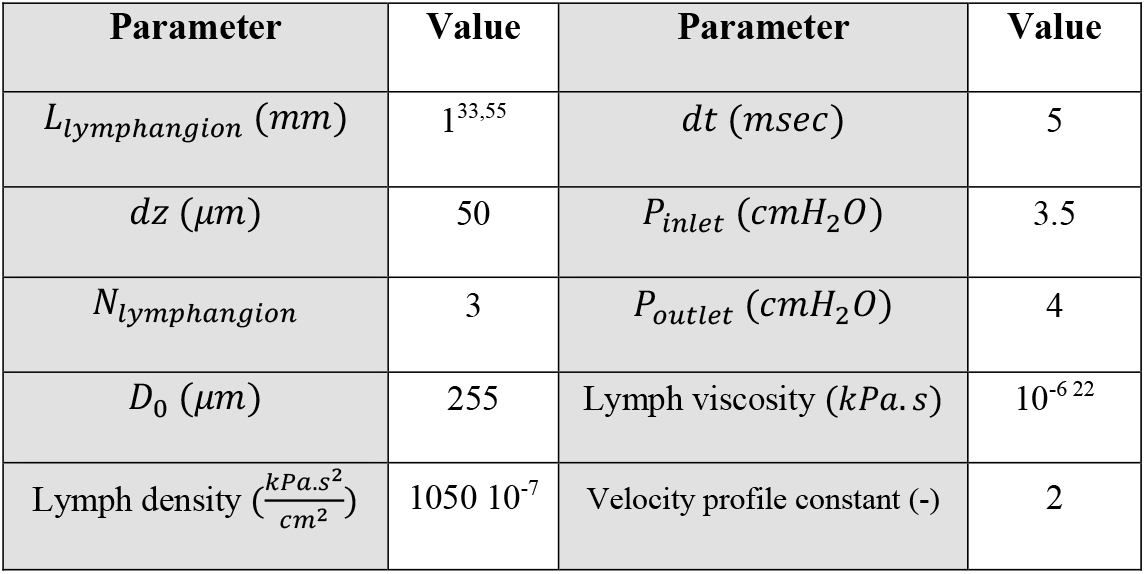
Simulation parameters

### Experimental data

To build and verify the model based on experimental data, we implement metrics from a paper by Gonzalez *et al*. that provides a useful set of contractility data for the healthy and the knock-out mouse strains^33^ (***Table 2***). The contraction propagation speed provided by Gonzalez are very close to what has been measured previously showing compliance with the literature^9,34^. The normal case is based on the wild-type (Cx45^fx/fx^) (Wt) mouse data and the abnormal cases are from Smmhc-CreER^T2^; Cx45^fx/fx^ [TMX] (2-4 weeks) (KO) mouse line. As was discussed, the effect of the direction of contraction wave propagation is an ongoing challenge^29,30^. To investigate that, three scenarios are considered for the KO model. In the first case, all contraction waves are antegrade while in the second case every other pacemaker generates a retrograde signal. In case three, all pacemaking signals are retrograde. The reason that these cases are not explored for the Wt model is that the contraction propagation velocity is high enough that the effect of direction of wave propagation is negligible.

**Table 2.** Contractility metrics used to prescribe the activation function

| Mouse line | Conduction Speed (cm/s) | Number of Initiation Sites (1/cm) | Contraction Frequency (1/min) | Contraction Period (sec) |
| --- | --- | --- | --- | --- |
| WT | 0.98 | 10 | 8.3 | 7.23 |
| KO | 0.07 | 31 | 15.8 | 3.80 |

## Results

### Activation function and lumen variations

As the input to the solver, the general activation function is generated based on the contractility metrics. ***Figure 6*** shows the time-space mapping of contractions during a cycle. By moving along the space domain, based on propagation speed and how close a node is from the pacemakers, the peak contraction arrives at different times. More quantitively, a parameter named activation integration (AI) is defined which is the volume integral under the space-time activation mapping. It turns out that AI is 189, 51, 69, and 49 for the Wt and the KO models, respectively. As AI correlates with the applied work on the vessel, we should expect a better pump efficiency in the Wt model. And although AI is close in the KO models, especially in Case 1 and 3, pumped lymph flow is different.

**Figure 6.**
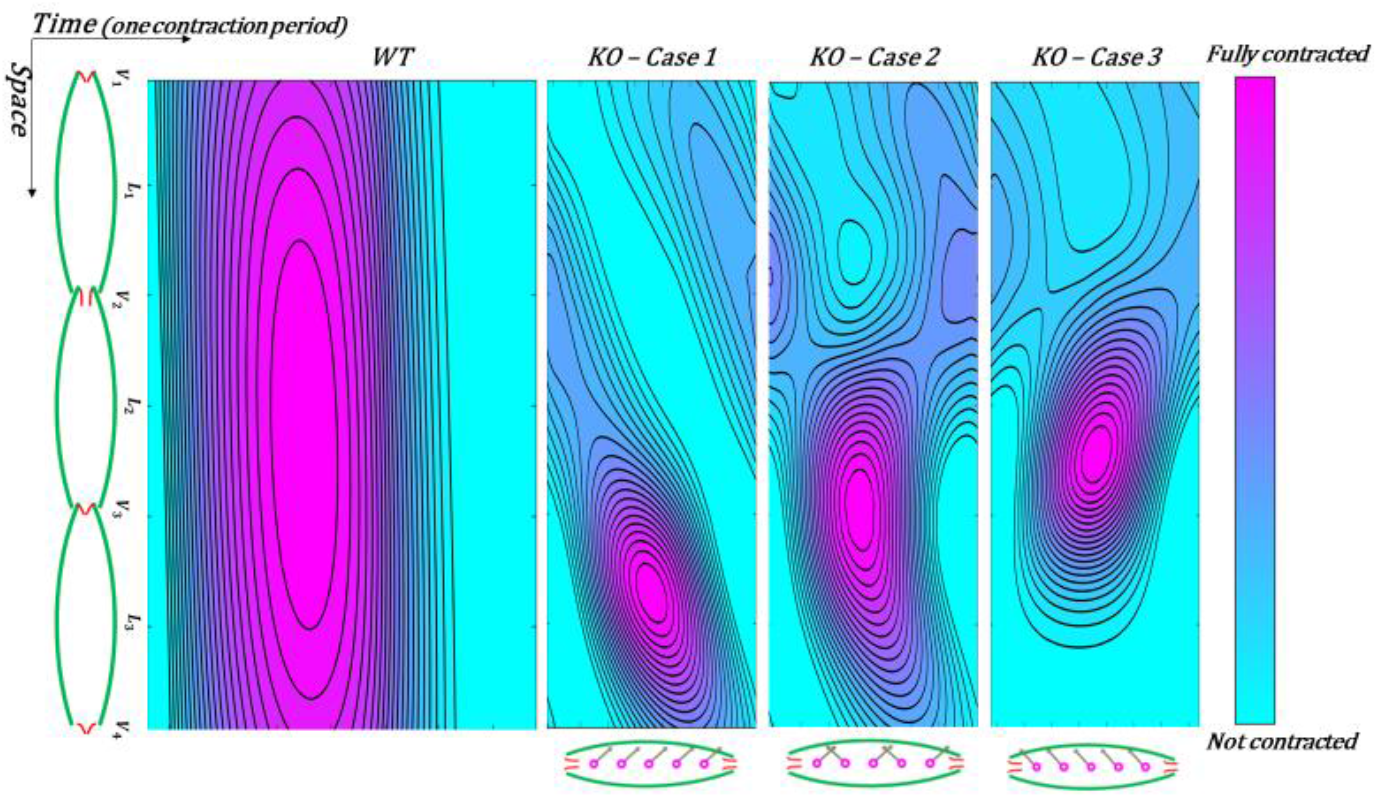
Heatmap of the general activaton function during a contraction cycle for Wt and KO models

The activation function is used in the model to calculate the radius and the compliance of each node. The resulting mapping of radius is demonstrated in ***Figure 7***. The radius patterns are essentially the opposite of the activation mappings, meaning that when the activation is zero, the vessel is relaxed and has the largest diameter and when it is fully contracted, the diameter is minimum.

**Figure 7.**
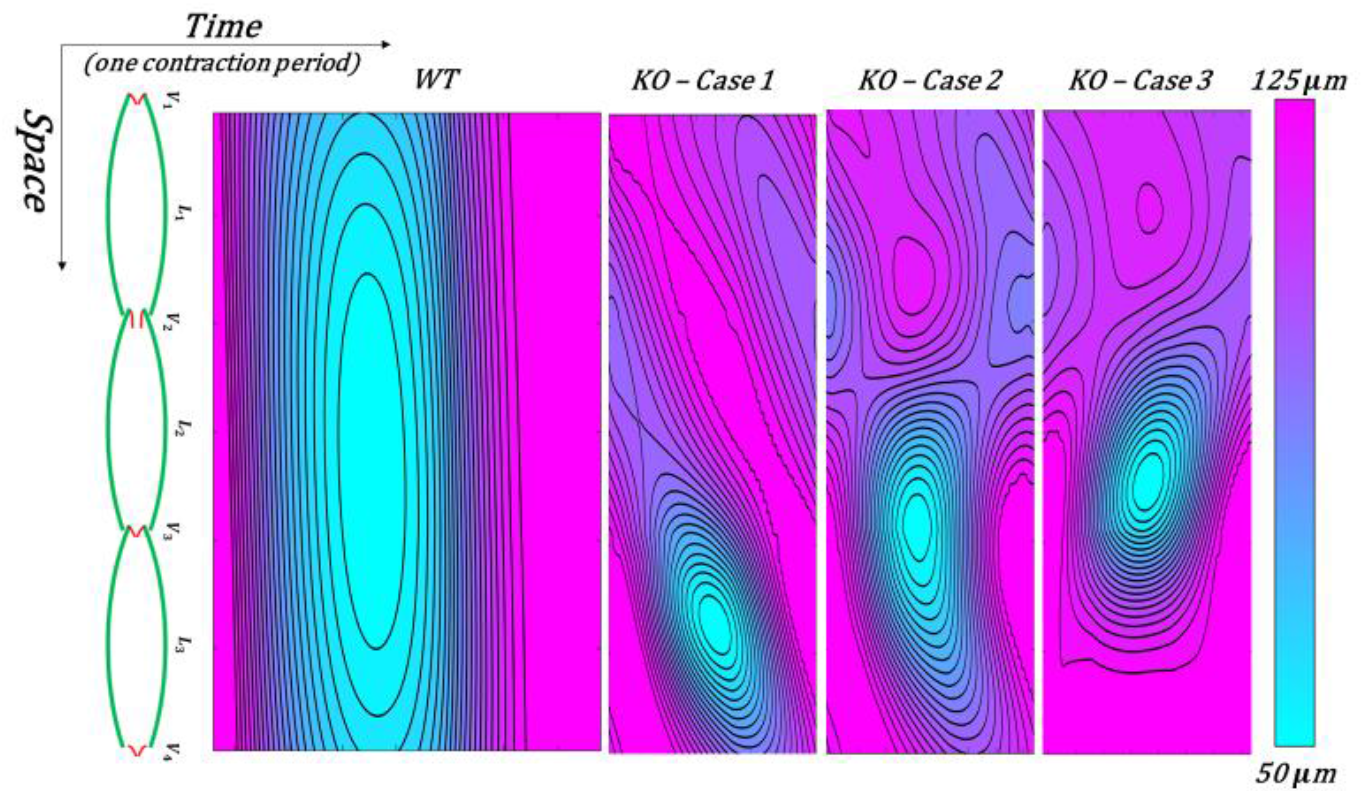
Heatmap of the radial contraction patterns during a contraction cycle

Another metric from the vessel mechanics is how the lengthwise average of lumen area changes in a contraction cycle. ***Figure 8*** shows the average lumen area contracts 80% in the Wt model followed by a remarkable relaxation period. But in all KO models, area reduction reaches at most 30%representing an inefficient pumping effect. Also, no relaxation period along the vessel is observed. This shows more engagement and fatigue the lymphatic muscle cells possibly undergo in abnormal contractility. To further expand on that, Ejection Fraction (EF) is defined based on the average end-diastolic (*A_EDA_*) and systolic (*A_ESA_*) areas (Equation (13)). EF shows how much lymph leaves the vessel with each contraction.

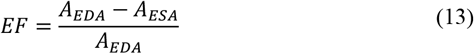

**Figure 8.**
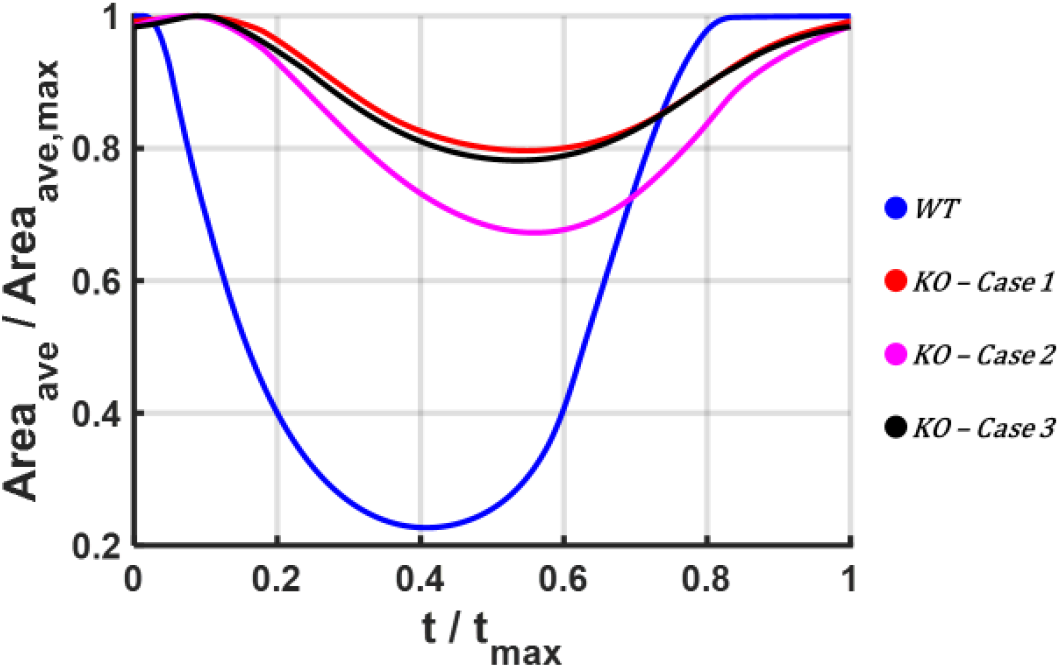
Normalized lengthwise average area vs. normalized contraction period

EF for the Wt model is about 0.78 but it varies between 0.2 and 0.33 in the KO models.

### Pressure patterns

***Figure 9b*** demonstrates the pressure profile of the middle node of each lymphangion during a contraction cycle. It is notable to see that in the Wt model, all lymphangions experience a pressure rise equivalent to the total pressure gradient of the chain. But in the KO models, only pressure in L3 bounces between *P_in_* and *P_out_*. ***Table 3*** shows the mean pressure in each lymphangion during a cycle. in the Wt model the mean pressure in all lymphangions stays around the average of *P_in_* and *P_out_*. But in the KO cases, only L3 is the working lymphangion that undergoes *P_in_* and *P_out_*, and L1 and L2 pressures stay near *P_in_*. This shows that in lymphedema some lymphangions may possibly experience a minimal pressure pulse during a contraction resulting in a further impaired contractility due to pressure-contraction couplings.

**Figure 9.**
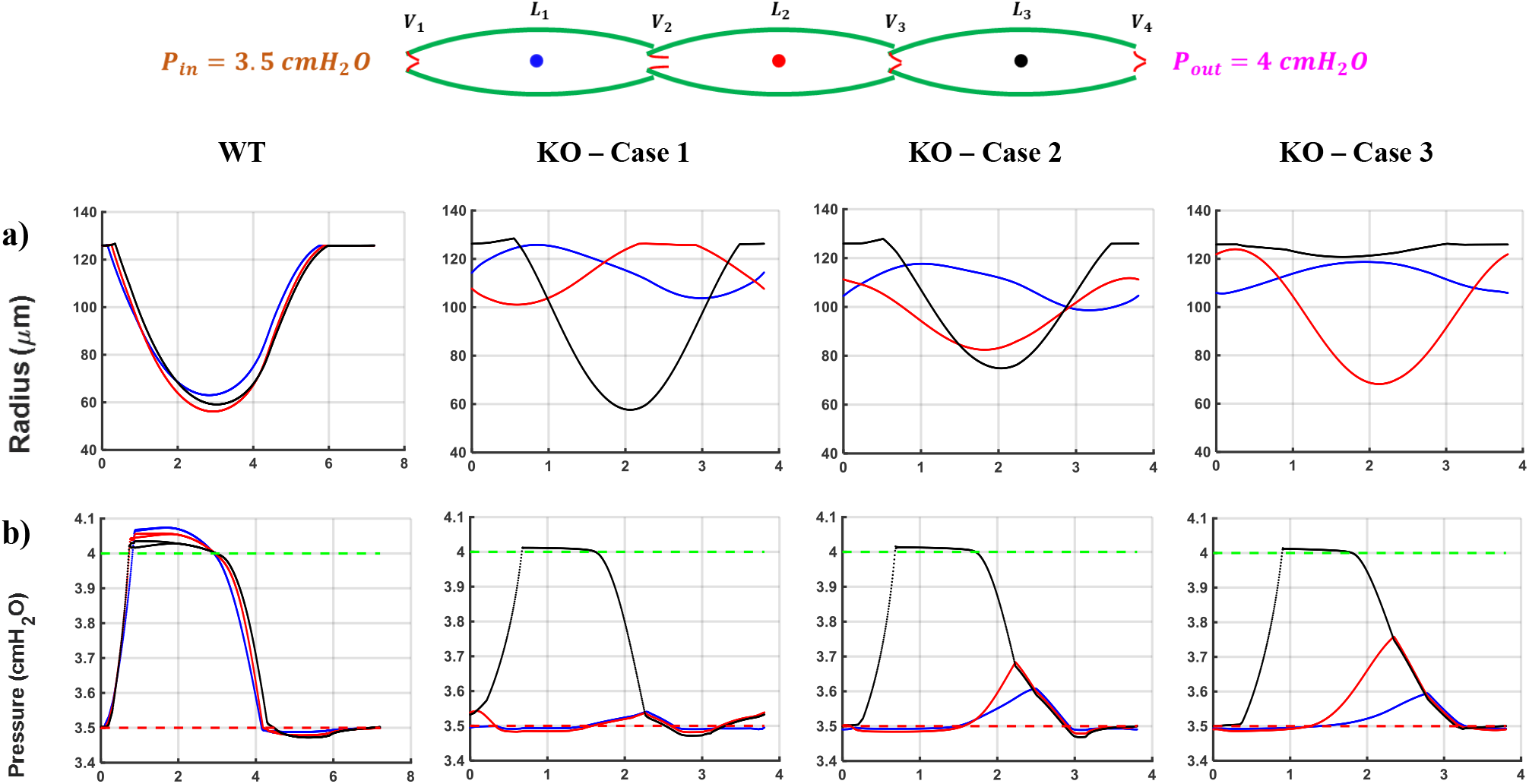
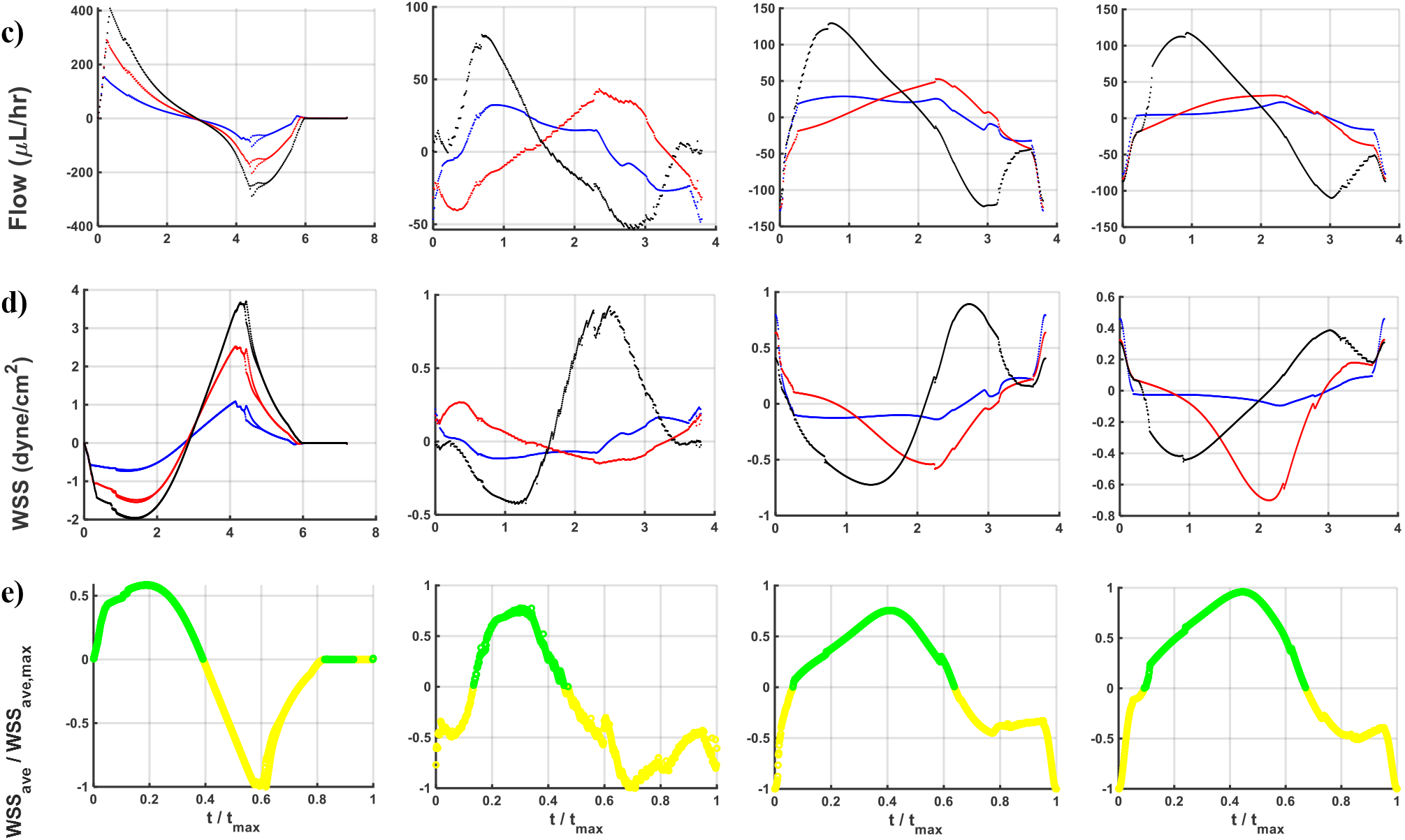
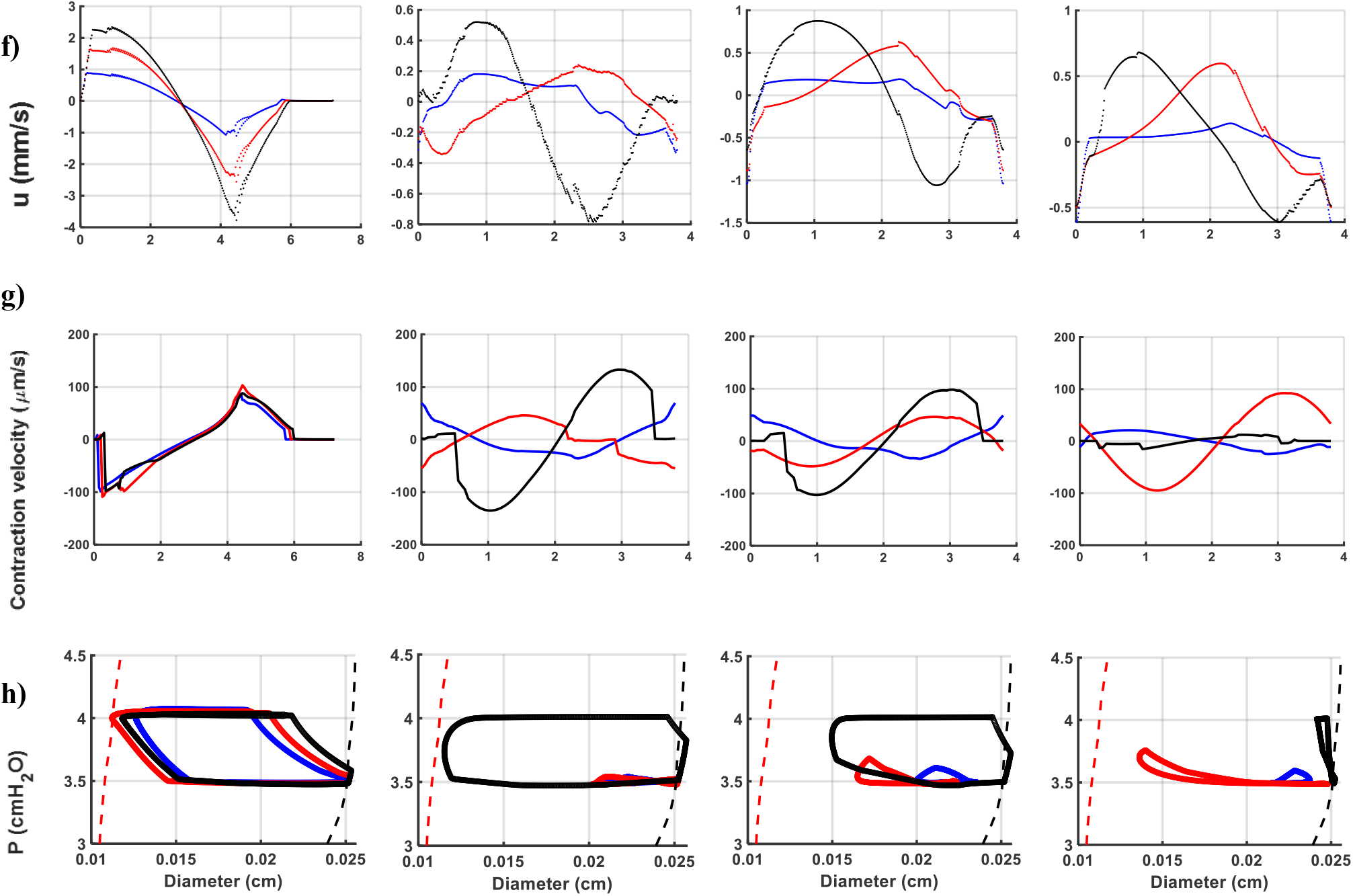
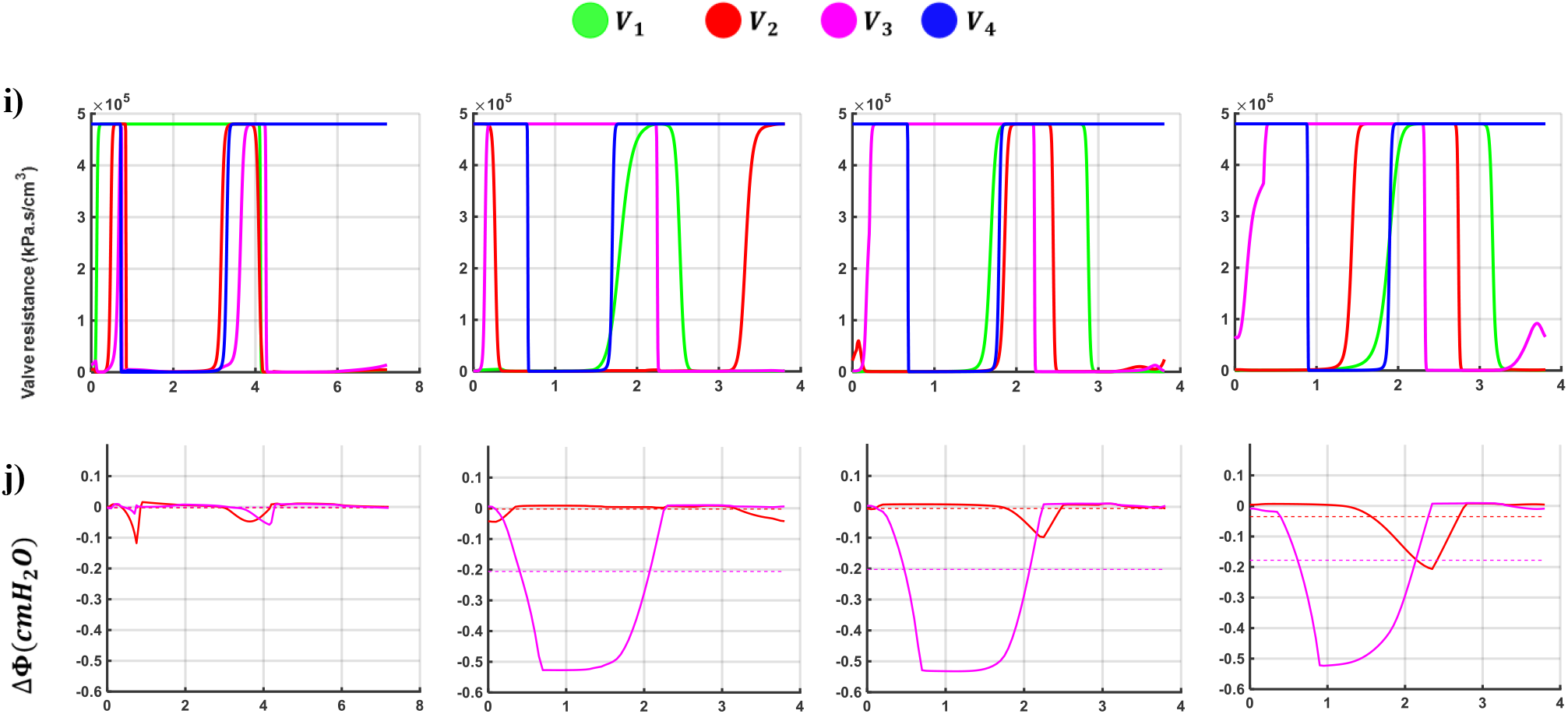
Modeling outcomes for WT and KO cases during a contraction cycle

**Table 3.** Mean pressure in each lymphangion during a contraction cycle

| Mean pressure ($cmH_2O$)<br>(Average of $P_{in}$ and $P_{out}$ is $3.75 cmH_2O$) | | | |
| --- | --- | --- | --- |
| Model | L1 | L2 | L3 |
| WT | 3.73 | 3.73 | 3.74 |
| KO – Case 1 | 3.50 | 3.50 | 3.71 |
| KO – Case 2 | 3.52 | 3.52 | 3.72 |
| KO – Case 3 | 3.52 | 3.55 | 3.73 |

### Flow patterns

***Figure 9c*** represents the flow patterns at the middle node of each lymphangion. A sudden impulse in flow concurrent to systole is seen in the Wt model which is followed by a dip due to relaxation and lumen expansion resulting in flow retracement. This gives a net pumped lymph flow of *5.44 μL/hr*. But a same behavior is not observed in the KO models. Although a relatively high flow passes the last lymphangion during systole, the first two lymphangions have opposing (zero or negative) flows. A similar behavior is seen during diastole where the flow in L3 retraces but L1 and L2 push the lymph forward. An analogy to this is pulling/pushing on the two ends of a syringe causing a hammer effect and sudden flow blockage causing flow blockage. The net cycle-averaged lymph flow is 2.9, *2.8* and *1.17 μL/hr* in Cases 1, 2, and 3 of the KO model, respectively.

The node with the maximum flow at each time step in a lymphangion is called the “Bottleneck” point. By looking at flow profiles along each lymphangion, we see that the bottleneck node stays stationary in all lymphangions in the WT model during a contraction cycle meaning that regardless of the phase (systolic or diastolic), a specific region has the highest lymph velocity. But this point is not stable in the KO models and it moves along the lymphangion. This could be an explanation of getting low net flow in the KO cases. In other words, at one moment point A has the maximum flow while point B located away from A has the maximum flow at a later time step resulting in an opposing lymph momentum. This shows that flow/pressure wave is destructed inside each lymphangion and the contraction energy is wasted locally and doesn’t help with the overall pump efficiency.

### Wall shear stress (WSS)

WSS is the next metric that is supposed to affect the contractility due to mechanosensitivity of lymphatic endothelial cells and their collaboration with the lymphatic muscle cells. We think that similar to blood vascular mechanics, both magnitude and smoothness of WSS affects the tone of contraction force. A challenging question is how WSS alters in lymphedema.

In the Wt model, the average, peak systolic and diastolic WSS are 0.08, −2, and 4 *dyn/cm*^2^, respectively (***Figure 9d***). In the KO models WSS varies between −1 to 1 *dyn/cm*^2^ and the average is 0.04, −0.07, and −0.07 *dyn/cm*^2^ in Cases 1, 2, and 3, respectively. The manifestation of the sudden impulse of pressure wave in the Wt model is again observed in the WSS patterns. Pushing lymph forward in systole causes a sudden drop in the WSS followed by a homogeneous reversal. This can be explained in two ways. First, the positive average WSS in the Wt model shows that along with the radial wall contractility effect on the lymph, the wall exerts a positive forward shear force which also helps further with the forward moving lymph. Also, the exerted shear stress on the wall is negative which is favorable in initiating the next contraction cycle. But this is not the case in the KO models where the average WSS is almost zero and the extremum WSS values are lower. All these show that both the extremum and the average wall shear stresses are highly disturbed in the KO models suggesting that a WSS-contractility coupling might be part of the inefficient pumping in lymphedema.

Along with magnitude, we postulate that smoothness of the WSS signal is important in the mechanotransduction events in the lymphatic muscle cells. To investigate that, we use a metric called signal-to-noise ratio (SNR) calculated from mean-squared error (MSE) of the WSS signal and it’s filtered form (Equation (14))^36^.

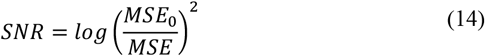

Where

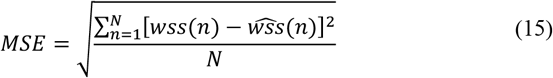

And

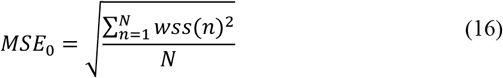

Where *h*(*n*) is the original signal containing noise, *N* is the length of the signal, and 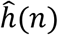 is the denoised signal calculated from sliding a window of length *windowSize* along the data, computing averages of the data contained in each window (Equation (17)).

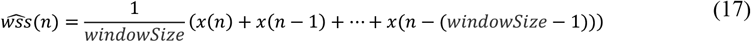

A huge gap between SNR of the Wt model (1.24) and the KO models (0.88, 0.69, and 0.70), shows how noisy the WSS signal is in the KO models. In other words, the LECs experience high shear disturbances that may disrupt the cellular signaling of the lymphatic muscle cells. So, both the magnitude and the SNR are favorable in the Wt model but they are highly disturbed in the KO models.

***Figure 9e*** shows the length-wise average of WSS during a contraction cycle; green represents positive and yellow is for negative values. Regardless of the cycle-average WSS values, these patterns show how symmetric *WSS_ave_* is during a contraction. The vessel wall experiences an identical shear stress during systole and diastole. But in the KO models, this identity is missing and the shear force is disorganized mostly during diastole. Also, sharp jumps in early systole and late diastole may exacerbate disruptions in the signaling mechanisms.

### Valve status

***Figure 9i*** shows the resistance of the valves during a contraction cycle. All valves operate in all modeling cases, but the order and the duration of openings/closures are different. In the Wt model, beginning of the contraction signal accompanies with sequentially closure of Valves 1 and 2 and opening of Valves 3,4 and then 2. Once the contraction decays and the pressure drops, the downstream valves close. In late diastole, V4 is the only closed valve while all other valves are open and ready for the next cycle. The biggest problem with the KO models is that during systole that we expect to only have the first valves closed to help with lymph propulsion, Valve 3 is closed as well showing that while some lymph is discharged from the last lymphangion, lymph in the first two lymphnsgions is trapped. Also, during diastole that we expect the downstream valves be closed and Valve 1 be open providing the lymph for the next cycle, Valves 2 and 3 are partially closed which means that less lymph volume can enter the vessel. So, only by looking at how valves operate, we can guess that the net pumped lymph is highly compromised in the KO models both because of less propelled lymph during systole and less stored lymph volume during diastole. It is understood that although all valves operate in the KO models, local pressure oppositions cause back flow in the valves and the net forward flow becomes negligible.

### Lymph velocity

***Figure 9.f*** shows that velocity patterns are similar to the WSS profiles. In the Wt model, a clear systolic peak velocity with the maximum velocity in the last lymphangion is followed by dipping of velocities with the maximum reverse velocity happening in the same location. This shows that the activation function amplifies the lymph momentum along the vessel, whether in propelling or absorbing lymph. But in the KO models, it is generally observed that the lymph velocity in the first two lymphangions partially oppose that of the last lymphangion both during systole and diastole. It is notable to see that in all KO models, the peak systolic and diastolic velocities do not exceed *1 mm/s*, while in the Wt model the peak velocity reaches up to 2.4 and −3.8 *mm/s*, respectively. Also, the velocity profiles are out-of-phase in the KO models and the systolic and diastolic phases of contraction are not distinguishable. Additionally, in the Wt model there is a resting state where all lymphangions are still and the net lymph flow is zero. But in all the KO models the contraction starts while a negative lymph velocity takes some of the contraction energy to overcome the backward momentum which means losing some of the pumping efficiency.

### Contraction velocity

Another metric is radial contraction velocity that is defined as the time derivative of radius (Equation (18)).

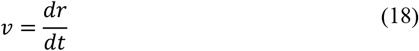

***Figure 9g*** shows that in the Wt model, all lymphangions contract almost simultaneously resulting in a contraction velocity of 100 *μm/s* followed by a similar positive velocity. It should be noted that the contraction happens in a short period of time (< 50ms) which shows that the vessel exerts a significant momentum to push the lymph. In Case 3 of the KO model, it is the middle lymphangion that has the highest contraction velocity, while in Case 1 and 2 the highest radial velocity happens in the last lymphangion. Interestingly, the peak systolic and diastolic velocities are around 100 *μm/s* in both the Wt and the KO models. In other words, although the vessel wall contracts with relatively high velocity in the KO models, it cannot generate enough propulsion force to move the lymph due to opposing pressure waves.

### Pressure-diameter (P-D) curves

As the most common representation tool, pressure-diameter curves show the mechanical behavior of a tissue (***Figure 9h***). The dashed lines belong to the pre- (red) and the peak-(black) twitch data. It is observed that in the Wt model, the pattern follows a clear pattern and bounces between the pre- and the peak-twitch curves meaning that all lymphangions experience both inlet and outlet pressures. So, we can suppose that in the Wt model, the total pressure gradient is experienced in each lymphangion during a contraction. But there are two major problems in the KO models. First, the patterns are not clearly identified and the diameter is mostly disturbed rather than getting smoothly contracted. Also, in the KO models each lymphangion stays closer whether to the inlet or outlet pressure and does not experience the total pressure gradient.

### Compliance-diameter (C-D) curves

Compliance is interpreted as how easy it is to inflate a vessel to a certain radial stretch and so it is inversely correlated to the stiffness. As in systole, the vessel wall gets contracted by the lymphatic muscle cells, we expect to see a decrease in compliance during systole. In other words, the lymphatic muscle cells become like the reinforced wires around the wall that strengthen the wall and doesn’t let the pressure increase the tube caliber but help push the lymph forward. ***Figure 10*** shows that there are three phases during a contraction. In early systole both diameter and compliance decrease followed by an iso-compliance phase where the diameter still decreases but the compliance is constant. Next, both the diameter and the compliance increase and return back to the pre-twitch states. The iso-compliance phase proves that in the healthy conditions, muscle cells twitch synchronically and stay activated while still the lumen closes. This doesn’t happen in the KO models, where the three phases are barely distinguishable and although the diameters decrease, the compliance doesn’t fluctuate significantly. In other words, regardless of how healthy the physiology of muscle cells is, the wall does not stiffen enough to propel the lymph.

**Figure 10.**
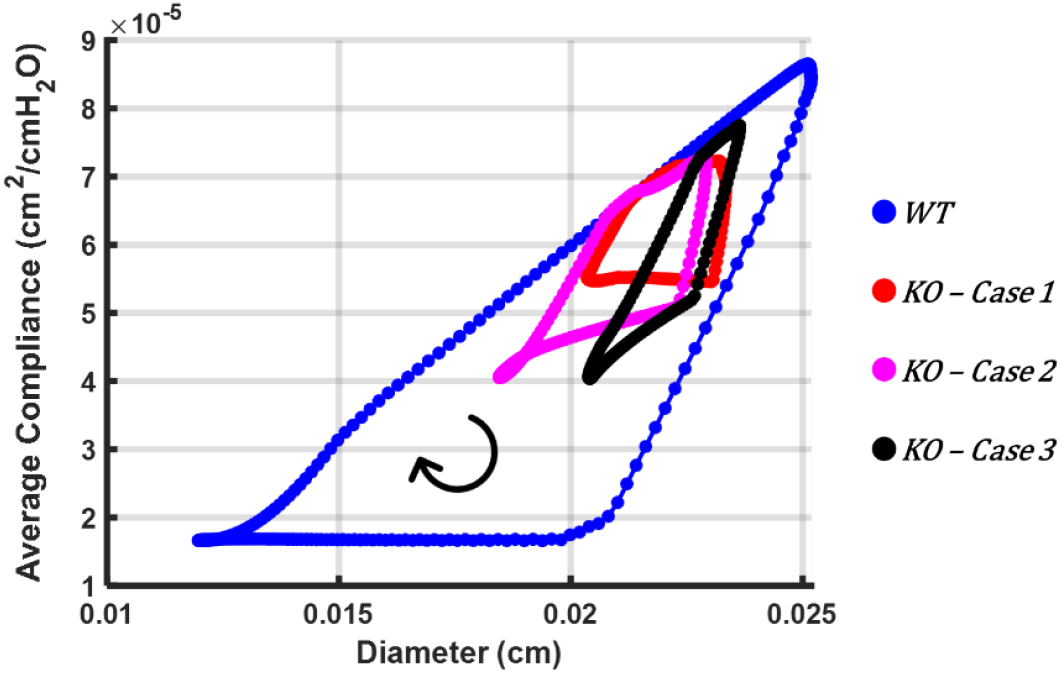
Average complaince vs. diameter of the lymphatic chain during a contraction cycle for the Wt and the KO case

### Energy loss at valves

A critical metric in studying valves in biomechanics is the loss of energy at valves due to leaflet performance and flow field. Energy loss is defined as the total energy difference at the valve. Total mechanical energy per volume (Φ) is defined in Equation (19)^37^.

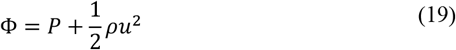

Where *P* is the luminal pressure, *ρ* is the lymph density, and *u* is the lymph velocity. The energy loss at a valve is the gap in the total mechanical energy (Equation (20)).

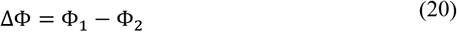

***Figure 9j*** shows the energy loss at the middle Valves 2 and 3. The average energy loss (dashed lines) is < 0.1 *cmH*_2_*O* in the Wt model, which stays the same for Valve 2 in the KO models. But it is −0.2 *cmH*_2_*O* in Valve 3 showing a remarkable energy loss. To better understand that, 0.2 *cmH*_2_*O* energy loss is equivalent to 40% of the total pressure head that the vessel pumps against. In other words, due to abnormal contractility, the vessel must pump harder to overcome both the total pressure gradient and the energy losses at the valves. That may explain the reason of inefficient pump action in KO vessels even in zero or favorable pressure gradients. Due to the pressure loss at the valves and probably flow disturbances on the leaflets, some remodeling mechanisms might trigger leading to changes in morphology of leaflets.

## Discussion

A 1-D mathematical model along with contractility metrics obtained from the experimental data of Wt and KO mouse lines were used to study the mechanical behavior of a lymphatic vessel under the two conditions^33^. We would expect different behaviors only by looking at time-space mapping of radius contractions (***Figure 7***). In terms of modeling outputs, luminal area reduction (***Figure 8***) and radius patterns (***Figure 9a***) demonstrate how deviated contractility metrics result in disturbed radius contractions and altered pump action. Reduction of the vessel caliber during systole in the Wt model (~50%) is similar to data measured in ex-vivo experiments^34^. More importantly, EF is 0.78 and [0.2-0.33] for the Wt and the KO models which mimic the data presented in the literature^33,38^. These observations show that using a spatiotemporal contraction distribution (Equations (10),(11)) and the weighted equation of radius (Equations (4)) and compliance (Equations (8)) are valid in simulating lymph mechanics.

In the Wt model the average and peak flow rates are 5.44 and 400 *μL/hr*, while they vary [1.17-2.9] and [80-150] *μL/hr* in the KO models. Dixon and Zawieja reported the average and peak flow rate of 14 *μL/hr* and 100 *μL/hr* ^38,39^. The former happens slightly in the Wt model and the latter attributes to the KO models which complies with the experimental observations.

These results become interesting when compared to data from wounded lymphatic vessels that were remodeled and demonstrated disturbed mechanical properties and contraction parameters^1^. If the KO model is comparable to the wounded case presented in their study, contractile frequency of the remodeled vessel for pressure gradient of 0.5 *cmH*_2_*O* is 2-3 times more than the control case, which is true in our model and the KO model mimics the decrease (~2.5 *μL/hr*) in driven lymph flow rate values calculated computationally with in-vivo data. So, by comparing the input (contraction frequency) and output (flow rate) of the model to their data, we can assume that the mapping used in this study for disease cases is relevant to a wounded/remodeled lymphatic vessel.

Dixon experimentally estimated the peak reversal lymph velocity of (−1 to 7) *mm/s* while the model estimated (−4 to −1) *mm/s*, showing a good compliance with the model^39^. Also, Zawieja estimated the peak lymph velocity between 2 and 9 *mm/s* which is almost observed in the model (0.5 and 2.5 mm/s)^38^. Most importantly, in an in-vivo study the peak forward lymph velocity reaches up to [2-3] *mm/s* which is very close to what has been calculated in the Wt model^10^. Also, Rahbar reported lymphocyte velocity and the average flow rate of [0.1-1.8] *mm/s* and [0-6] *μL/hr* which includes all velocities and flows from both the Wt and the KO models^40^. Interestingly, the peak diastolic negative velocity is higher than the systolic positive velocity in all models, while the net pumped flow is still positive. This shows the importance of measuring the area as well as the velocity profile to calculate the flow rate.

WSS values from the model are partially comparable to values reported by Mukherjee, especially by considering the difference in diameters^41^. They showed that the mean WSS is approximately 0.6 *dyn/cm*^2^ with the peak of 8 *dyn/cm*^2^ while they are 0.17 and 10-20 *dyn/cm*^2^ in the model. Also, Zawieja showed that mean WSS is [0.4-0.6] *dyn/cm*^2^ with the peak of [3-10] *dyn/cm^2^*, which are close to the model calculations^38^. Also, Wilson and Dixon estimated the average WSS of 0.64, Kornuta reported [0-2], and Rahbar stated 0.15 *dyn/cm*^2^ in control cases which happens in the Wt model^14,39,40,42^. So, in terms of WSS, we can postulate that the model calculation falls into the range of wall shear stresses that were reported in the literature. Also, we should notice that the average WSS is two orders of magnitude smaller than the peak systolic and diastolic values, showing that any inaccuracy or noise in measuring WSS with experimental methods is highly prune to erroneous readings.

P-D patterns are similar to experiments in terms of sequence and amount of diameter reductions and pressure rises^43^. And chaotic P-D patterns in the KO models can be used as a visual tool in showing abnormal contractility of lymphatic vessels. Also, P-D patterns in the Wt model prove that using the weighted average of diameters along with an activation parameter is a good substitution to the constitutive equation.

Disparities between KO models prove that the directionality of pacemaking signals highly affects the pump efficiency which has been addressed before^30^ and implementing such pacemaking behavior is not feasible with lumped parameter models. By using contraction mappings obtained from imaging data, we can easily modify Gaussian functions to accurately prescribe contractions of a lymphatic vessel and estimate pressure/flow patterns.

Finally, by using a 1-D formulation of the lymphatic network and the spatiotemporal contraction wave, we provide an experimentally motivated simulation of lymphatic mechanics and the effects of impaired contractility on pump efficacy due to Cx45 knocked out. The model further provides estimates of mechanical parameters (pressure, flow, wall shear stress) that are difficult to measure experimentally, and can be used to perform parametric studies and explore the role of mechanical properties (e.g., stiffness, contraction propagation, valve resistance) on lymphatic function and the progression of lymphedema.

### Limitations and Future Works

To expand the current model, more studies relevant to lymphedema could be carried out such as: abnormal wall mechanics^1^, leakiness of the secondary valves^44^, abnormal initial lymphatic vessels, and leakiness of the primary valves^22^. Among these, adding initial lymphatic vessels and considering leakiness of the primary valves could provide a better picture of how lymphedema develops. Also, in this model the contractions are decoupled from the pressure and WSS effects, which is not the case in lymphatics^15,17,40,41,45,46^. So, to add another layer of complexity, one may add pressure/WWS and contraction couplings to include the mechanical effects. Or these couplings could be implemented to study injury and remodeling^1,47^. To further expand the current model, we may look at the effect of axial pressure gradient on pump efficiency and correlation of number of lymphangions and the pressure gradient^48,49^. Also, the effect of contractility on passive (conduit) and active (pump) lymph transport could be addressed with this 1-D model^50^. Executing a parametric study similar to ^51^ could be another interesting topic to investigate the contribution of each parameter in the development of lymphedema. Also, by using a correction factor proposed by Razavi *et al*. we can easily scale the pre- and peak-twitch curves to prescribe the mechanical response under disturbed chemical (nitric oxide) environment^45^. Finally, due to high energy loss at the valves in KO models, the model suggests to study the morphology of valve leaflets in lymphedema and how they possibly exacerbate the pumping efficiency.

## Conclusions

In summary, we have developed a mathematical model based on a 1-D framework of the mechanical balance laws and experimental data of lymphatic contractility. Each set of contractility data enforces a pattern of contractions leading to different lymphatic mechanics. Due to deviation in pacemaking parameters, contractions are non-harmonic and non-sequential in the knock-out models. Although diameters contract significantly, opposing contractions and pressure barriers result in a negligible pumped flow in the KO models. Similarly, abnormal WSS may address abnormal contractility in lymphedema in long-term periods. Disturbed contraction velocity in the KO mouse models show that the work done on the vessel wall is not used as the energy source to push the flow and it is wasted locally. And, although the valves open and close in all models, the net lymph flow is zero in the KO models. It should be reminded that in this model, we only focus on contractions, and the wall mechanics and valve behavior are assumed to be unchanged in the disease cases. Finally, the current model could be a useful tool in studying the lymph transport in conjunction with an imaging technique. In other words, contractile metrics obtained from online time-space radius mappings can be used to estimate mechanical metrics (flow, velocity, WSS, etc.) of a lymphatic specimen. Although the experimental data have been used to build the model, still there is a pressing need to extensively quantify the lymph velocity profile non-invasively whether by using NIR imaging, DOCT techniques, or other methods^52,53^. The current study enables us to easily apply changes in pacemaking parameters based on the type and mechanics of a lymphatic vessel^54^. Also, the model proves that only by analyzing some aspects of the mechanical behavior of a lymphatic vessel, we cannot get a clear picture of the pump efficiency. For example, we saw that the peak WSS, contraction velocities, or even radius reductions are similar in both the Wt and the KO models, but the resulting driven lymph flows are highly deviated in the KO models. So, lymphatic contractility is an extensive problem that necessitates studying all aspects simultaneously.

