## Appendix A for "A 1D Model Characterizing the Role of Spatiotemporal Contraction Distributions on Lymph Transport"

***Linearization and discretization of collecting vessel equations***

Starting from balance of mass and linear momentum

$$C\frac{\partial P}{\partial t}+\frac{\partial Q}{\partial z}=0 \quad (A1)$$

$$\frac{\rho}{A}\frac{\partial Q}{\partial t}+\frac{\partial P}{dz}=\hat{h} \quad (A2)$$

where $\hat{h}$ denotes values taken from a previous time-point and

$$\hat{h}=\frac{f}{A}-\frac{\rho}{A}\frac{\partial}{\partial z}\left(\frac{Q^2}{A}\right) \quad (A3)$$

To solve equations, we discretize the lymphatic domain into a number of nodes that define a discrete number of non-overlapping two-node elements. Across any set of elements,

$$\int(\blacksquare)dz=\sum_{e=1}^{N_e}\int_e(\blacksquare)dz \quad (A4)$$

where $N_e$ is the number of elements over the section of interest. We employ the trapezoid rule to spatially integrate over z. Consider an arbitrary 1D element, $e_j$, from node $j$ to node $j+1$. The integration over this element, for each term in equations is given as,

$$\int_{e_j}\left(C\frac{\partial P}{\partial t}\right)dz\approx\left(C_j\frac{\partial P_j}{\partial t}+C_{j+1}\frac{\partial P_{j+1}}{\partial t}\right)\frac{\Delta z_{e_j}}{2} \quad (A5)$$

$$\int_{e_j}\frac{\partial P}{\partial z}dz\approx P_{j+1}-P_j \quad (A6)$$

$$\int_{e_j}\left(\frac{\rho}{A}\frac{\partial Q}{\partial t}\right)dz\approx\left(\frac{\rho}{A_j}\frac{\partial Q_i}{\partial t}+\frac{\rho}{A_{j+1}}\frac{\partial Q_{j+1}}{\partial t}\right)\frac{\Delta z_{e_j}}{2} \quad (A7)$$

$$\int_{e_j}\frac{\partial Q}{\partial z}dz\approx Q_{j+1}-Q_j \quad (A8)$$

$$\int_{e_j} h dz \approx \left(h_j + h_{j+1}\right) \frac{\Delta z_{e_j}}{2} \qquad (A9)$$

Time derivatives are approximated using a second-order backward difference scheme; e.g.,

$$\frac{\partial P_i}{\partial t} = \frac{3P_i^{t+\Delta t} - 4P_i^t + P_i^{t-\Delta t}}{2\Delta t} \qquad (A10)$$

For $j = 1$ to $N_z - 1$ and $j = N_z$, respectively, we approximate the spatial derivatives as

$$\frac{\partial}{\partial z}\left(\frac{{Q_j^i}^2}{A_j^i}\right) \cong \left(\frac{{Q_{j+1}^i}^2}{A_{j+1}^i} - \frac{{Q_j^i}^2}{A_j^i}\right)\frac{1}{\Delta z_{e_j}} \qquad (A11)$$

$$\frac{\partial}{\partial z}\left(\frac{{Q_{N_z}^i}^2}{A_{N_z}^i}\right) \cong \left(\frac{{Q_{N_z}^i}^2}{A_{N_z}^i} - \frac{{Q_{N_z-1}^i}^2}{A_{N_z-1}^i}\right)\frac{1}{\Delta z_{e_{N_z}}} \qquad (A12)$$

where $j = N_z$ is the last node of the vessel.

Substituting these equations into equations (A1,2) we get

$$\left(C_j^i \frac{3P_j^{i+1} - 4P_j^i + P_j^{i-1}}{2\Delta t} + C_{j+1}^i \frac{3P_{j+1}^{i+1} - 4P_{j+1}^i + P_{j+1}^{i-1}}{2\Delta t}\right)\frac{\Delta z_{e_j}}{2} + Q_{j+1}^{i+1} - Q_j^{i+1} = 0 \qquad (A13)$$

$$\left(\frac{\rho}{A_j^i}\frac{3Q_j^{i+1} - 4Q_j^i + Q_j^{i-1}}{2\Delta t} + \frac{\rho}{A_{j+1}^i}\frac{3Q_{j+1}^{i+1} - 4Q_{j+1}^i + Q_{j+1}^{i-1}}{2\Delta t}\right)\frac{\Delta z_{e_j}}{2} + P_{j+1}^{i+1} - P_j^{i+1} = \left(h_j + h_{j+1}\right)\frac{\Delta z_{e_j}}{2} \qquad (A14)$$

Equations (A13,14) may be re-written in matrix form as,

$$F_{e_j} M_{e_j}^{i+1} = G_{e_j} N_{e_j}^{i+1} + h_{e_j} \qquad (A15)$$

where, with the flow boundary condition, it is useful to let $M_{e_j}^{i+1}$ be defined in terms of the nodal pressures and $N_{e_j}^{i+1}$ be in terms of the nodal flows.

The initiating node and internal nodes $e_1$ to $e_{N_z-1}$ may be written:

$$\left[M_{e_j}^{i+1}\right] = \begin{bmatrix} P_j^{i+1} \\ P_{j+1}^{i+1} \end{bmatrix} \qquad (A16)$$

$$\left[N_{e_j}^{i+1}\right] = \begin{bmatrix} Q_j^{i+1} \\ -Q_{j+1}^{i+1} \end{bmatrix} \tag{A17}$$

By re-organizing equations A13,14:

$$C_j^i \frac{3\Delta z_{e_j}}{4\Delta t} P_j^{i+1} + C_{j+1}^i \frac{3\Delta z_{e_j}}{4\Delta t} P_{j+1}^{i+1} = Q_j^{i+1} + (-Q_{j+1}^{i+1}) + C_j^i \frac{\Delta z_{e_j}}{4\Delta t}\left(4P_j^i - P_j^{i-1}\right) + C_{j+1}^i \frac{\Delta z_{e_j}}{4\Delta t}\left(4P_{j+1}^i - P_{j+1}^{i-1}\right) \tag{A18}$$

$$-P_j^{i+1} + P_{j+1}^{i+1} = -\frac{\rho}{A_j^i}\frac{3\Delta z_{e_j}}{4\Delta t} Q_j^{i+1} + \frac{\rho}{A_{j+1}^i}\frac{3\Delta z_{e_j}}{4\Delta t}(-Q_{j+1}^{i+1}) + \left(h_j + h_{j+1}\right)\frac{\Delta z_{e_j}}{2} + \frac{\rho}{A_j^i}\frac{\Delta z_{e_j}}{4\Delta t}\left(4Q_j^i - Q_j^{i-1}\right) + \frac{\rho}{A_{j+1}^i}\frac{\Delta z_{e_j}}{4\Delta t}\left(4Q_{j+1}^i - Q_{j+1}^{i-1}\right) \tag{A19}$$

So the matrices are,

$$[F_{e1}] = \begin{bmatrix} C_j^i \frac{3\Delta z_{e_j}}{4\Delta t} & C_{j+1}^i \frac{3\Delta z_{e_j}}{4\Delta t} \\ -1 & 1 \end{bmatrix} \tag{A20}$$

$$[G_{e1}] = \begin{bmatrix} 1 & 1 \\ -\frac{\rho}{A_j^i}\frac{3\Delta z_{e_j}}{4\Delta t} & \frac{\rho}{A_{j+1}^i}\frac{3\Delta z_{e_j}}{4\Delta t} \end{bmatrix} \tag{A21}$$

$$[h_{e1}] = \begin{bmatrix} C_j^i \frac{\Delta z_{e_j}}{4\Delta t}\left(4P_j^i - P_j^{i-1}\right) + C_{j+1}^i \frac{\Delta z_{e_j}}{4\Delta t}\left(4P_{j+1}^i - P_{j+1}^{i-1}\right) \\ \left(h_j + h_{j+1}\right)\frac{\Delta z_{e_j}}{2} + \frac{\rho}{A_j^i}\frac{\Delta z_{e_j}}{4\Delta t}\left(4Q_j^i - Q_j^{i-1}\right) + \frac{\rho}{A_{j+1}^i}\frac{\Delta z_{e_j}}{4\Delta t}\left(4Q_{j+1}^i - Q_{j+1}^{i-1}\right) \end{bmatrix} \tag{A22}$$

For the last element we have,

$$\left[M_{e_j}^{i+1}\right] = \begin{bmatrix} P_j^{i+1} \\ Q_{j+1}^{i+1} \end{bmatrix} \tag{A23}$$

$$\left[N_{e_j}^{i+1}\right] = \begin{bmatrix} Q_j^{i+1} \\ P_{j+1}^{i+1} \end{bmatrix} \tag{A24}$$

So,

$$C_j^i \frac{3\Delta z_{e_j}}{4\Delta t} P_j^{i+1} + Q_{j+1}^{i+1} = Q_j^{i+1} - C_{j+1}^i \frac{3\Delta z_{e_j}}{4\Delta t} P_{j+1}^{i+1} + C_j^i \frac{\Delta z_{e_j}}{4\Delta t}\left(4P_j^i - P_j^{i-1}\right) + C_{j+1}^i \frac{\Delta z_{e_j}}{4\Delta t}\left(4P_{j+1}^i - P_{j+1}^{i-1}\right) \tag{A25}$$

$$-P_j^{i+1} + \frac{\rho}{A_{j+1}^i}\frac{3\Delta z_{e_j}}{4\Delta t} Q_{j+1}^{i+1} = -\frac{\rho}{A_j^i}\frac{3\Delta z_{e_j}}{4\Delta t} Q_j^{i+1} - P_{j+1}^{i+1} + \left(h_j + h_{j+1}\right)\frac{\Delta z_{e_j}}{2} + \frac{\rho}{A_j^i}\frac{\Delta z_{e_j}}{4\Delta t}\left(4Q_j^i - Q_j^{i-1}\right) + \frac{\rho}{A_{j+1}^i}\frac{\Delta z_{e_j}}{4\Delta t}\left(4Q_{j+1}^i - Q_{j+1}^{i-1}\right) \tag{A26}$$

And the matrices are,

$$[F_{e1}] = \begin{bmatrix} C_j^i \dfrac{3\Delta z_{e_j}}{4\Delta t} & 1 \\ -1 & \dfrac{\rho}{A_{j+1}^i} \dfrac{3\Delta z_{e_j}}{4\Delta t} \end{bmatrix} \qquad (A27)$$

$$[G_{e1}] = \begin{bmatrix} 1 & -C_{j+1}^i \dfrac{3\Delta z_{e_j}}{4\Delta t} \\ -\dfrac{\rho}{A_j^i} \dfrac{3\Delta z_{e_j}}{4\Delta t} & -1 \end{bmatrix} \qquad (A28)$$

$$[h_{e1}] = \begin{bmatrix} C_j^i \dfrac{\Delta z_{e_j}}{4\Delta t}\left(4P_j^i - P_j^{i-1}\right) + C_{j+1}^i \dfrac{\Delta z_{e_j}}{4\Delta t}\left(4P_{j+1}^i - P_{j+1}^{i-1}\right) \\ \left(h_j + h_{j+1}\right)\dfrac{\Delta z_{e_j}}{2} + \dfrac{\rho}{A_j^i}\dfrac{\Delta z_{e_j}}{4\Delta t}\left(4Q_j^i - Q_j^{i-1}\right) + \dfrac{\rho}{A_{j+1}^i}\dfrac{\Delta z_{e_j}}{4\Delta t}\left(4Q_{j+1}^i - Q_{j+1}^{i-1}\right) \end{bmatrix} \qquad (A29)$$

***Discretization of secondary valve equations***

Starting from conservation of mass of the secondary valves, we assume that the valves are compliant meaning the volume changes in time,

$$\frac{\partial A}{\partial t} + \frac{\partial Q}{\partial z} = 0 \qquad (A30)$$

We employ the trapezoid rule to spatially integrate over z. Consider an arbitrary 1D element, $e_j$, from node $j$ to node $j + 1$,

$$\int_{e_j} \left(\frac{\partial A}{\partial t}\right) dz \approx \left(\frac{\partial A_j}{\partial t} + \frac{\partial A_{j+1}}{\partial t}\right)\frac{\Delta z_{e_j}}{2} \qquad (A31)$$

Time derivatives are approximated using a second-order backward difference scheme; e.g.,

$$\frac{\partial A_j}{\partial t} = \frac{3A_j^{t+\Delta t} - 4A_j^t + A_j^{t-\Delta t}}{2\Delta t} \qquad (A32)$$

So,

$$\int_{e_j} \left(\frac{\partial A}{\partial t}\right) dz \approx \left(\frac{3A_j^{t+\Delta t} - 4A_j^t + A_j^{t-\Delta t}}{2\Delta t} + \frac{3A_{j+1}^{t+\Delta t} - 4A_{j+1}^t + A_{j+1}^{t-\Delta t}}{2\Delta t}\right)\frac{\Delta z_{e_j}}{2} \qquad (A33)$$

Also,

$$\int_{e_j} \left(\frac{\partial Q}{\partial z}\right) dz \approx Q_{j+1} - Q_j \tag{A34}$$

So,

$$\left(\frac{3A_j^{t+\Delta t} - 4A_j^t + A_j^{t-\Delta t}}{2\Delta t} + \frac{3A_{j+1}^{t+\Delta t} - 4A_{j+1}^t + A_{j+1}^{t-\Delta t}}{2\Delta t}\right)\frac{\Delta z_{e_j}}{2} + Q_{j+1} - Q_j = 0 \tag{A35}$$

Or,

$$0 = -Q_j + Q_{j+1} + \left(\frac{3A_j^{t+\Delta t} - 4A_j^t + A_j^{t-\Delta t}}{2\Delta t} + \frac{3A_{j+1}^{t+\Delta t} - 4A_{j+1}^t + A_{j+1}^{t-\Delta t}}{2\Delta t}\right)\frac{\Delta z_{e_j}}{2} \tag{A36}$$

Also, for the secondary valves,

$$Q_j^{i+1} = \frac{P_j^{i+1} - P_{j+1}^{i+1}}{R_v^i} \tag{A37}$$

or

$$P_j^{i+1} - P_{j+1}^{i+1} = R_v^i Q_j^{i+1} \tag{A38}$$

$R_v^i$ which is the resistance of valve $i$, is a function of pressure difference at the valve.

To re-write the equations in matrix form we have,

$$F_{e_j} M_{e_j}^{i+1} = G_{e_j} N_{e_j}^{i+1} + h_{e_j} \tag{A39}$$

where

$$\left[M_{e_j}^{i+1}\right] = \begin{bmatrix} P_j^{i+1} \\ P_{j+1}^{i+1} \end{bmatrix} \tag{A40}$$

$$\left[N_{e_j}^{i+1}\right] = \begin{bmatrix} Q_j^{i+1} \\ -Q_{j+1}^{i+1} \end{bmatrix} \tag{A41}$$

$$[F_{e1}] = \begin{bmatrix} 0 & 0 \\ 1 & -1 \end{bmatrix} \tag{A42}$$

$$[G_{e1}] = \begin{bmatrix} -1 & -1 \\ R_v^i & 0 \end{bmatrix} \quad (A43)$$

$$[h_{e1}] = \begin{bmatrix} \left( \dfrac{3A_j^{t+\Delta t} - 4A_j^t + A_j^{t-\Delta t}}{2\Delta t} + \dfrac{3A_{j+1}^{t+\Delta t} - 4A_{j+1}^t + A_{j+1}^{t-\Delta t}}{2\Delta t} \right) \dfrac{\Delta z_{e_j}}{2} \\ 0 \end{bmatrix} \quad (A44)$$

After discretizing all the equations including (1-D formulation and secondary valve), we need to assemble all the equations into a matrix of coefficients and unknows and then by solving for the unknowns, the pressures and flows are calculated.
